# An Engineered Amber-Emitting Nano Luciferase and Its Use for Immunobioluminescence Imaging in Vivo

**DOI:** 10.1101/2022.05.23.493143

**Authors:** Ying Xiong, Yiyu Zhang, Zefan Li, Md Shamim Reza, Xinyu Li, Xiaodong Tian, Huiwang Ai

**Author notes:** State Key Laboratory of Reproductive Regulation and Breeding of Grassland Livestock, School of Life Sciences, Inner Mongolia University, Hohhot, 010070, China (Xinyu Li).

## Abstract

The NanoLuc luciferase (NLuc) and its furimazine (FRZ) substrate have revolutionized bioluminescence (BL) assays and imaging. However, the use of the NLuc-FRZ luciferase-luciferin pair for mammalian tissue imaging is hindered by the low tissue penetration of the emitting blue photons. Here, we present the development of an NLuc mutant, QLuc, which catalyzes the oxidation of a synthetic QTZ luciferin for bright and red-shifted emission peaking at ∼ 585 nm. This amber-light-emitting luciferase-luciferin pair exhibited improved performance for imaging deep-tissue targets in live mice. Leveraging this novel bioluminescent reporter, we further pursued *in vivo* immunobioluminescence imaging (immunoBLI), which used a fusion protein of a single-chain variable antibody fragment (scFv) and QLuc for molecular imaging of tumor-associated antigens in a xenograft mouse model. As one of the most red-shifted NLuc variants, we expect QLuc to find broad applications in noninvasive imaging in mammals. Moreover, the immunoBLI method complements immunofluorescence imaging and immuno-positron emission tomography (immunoPET), serving as a convenient and nonradioactive molecular imaging tool for animal models in basic and preclinical research.

## INTRODUCTION

BL imaging (BLI), one of the most sensitive noninvasive imaging modalities, has been widely utilized in basic and preclinical studies.^1,2^ Commonly used bioluminescent reporters belong to one of the two categories: luciferases derived from insects, such as firefly luciferase (FLuc), and luciferases derived from marine creatures, such as *Renilla* luciferase (RLuc).^3-5^ Insect luciferases typically use D-luciferin as the substrate, while marine luciferases usually oxidize the coelenterazine (CTZ) luciferin for bioluminescence (**Figure 1**). Historically, researchers preferred insect luciferases since D-luciferin is chemically stable and accessible. Also, compared to marine luciferases, insect luciferases emit longer-wavelength light, which interacts less with mammalian tissue, resulting in increased tissue penetration for imaging. However, insect luciferases consume adenosine triphosphate (ATP) for D-luciferin activation and photon emission, so they only function properly when sufficient ATP is present.^6^

**Figure 1.**
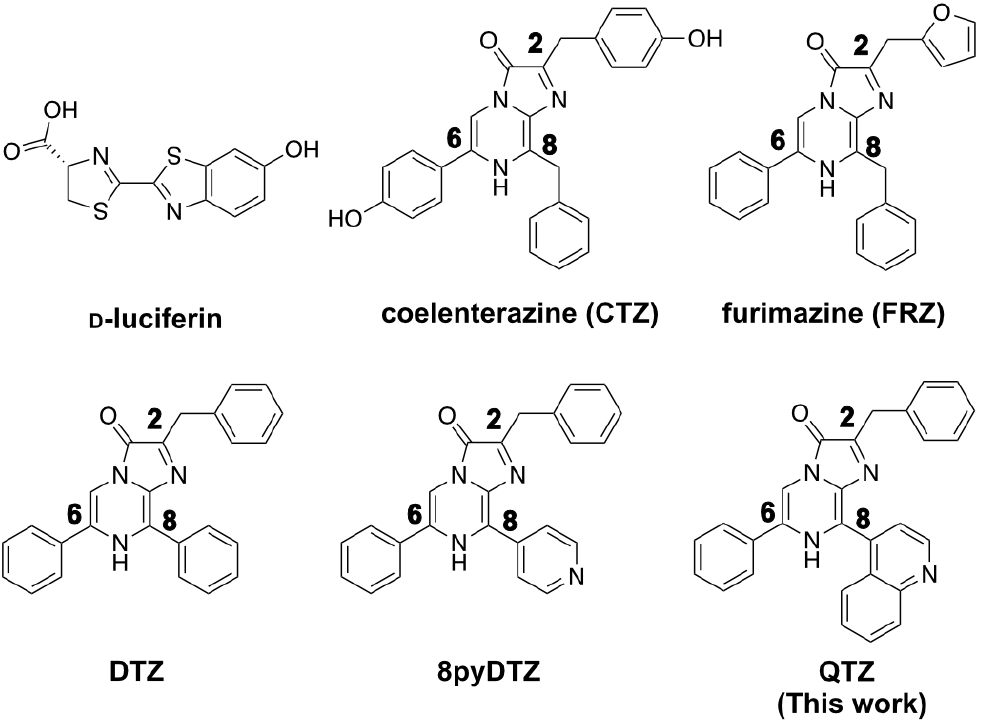
Chemical structures of D-luciferin, coelenterazine (CTZ), and several representative CTZ analogs. The CTZ analogs are labeled for the C2, C6, and C8 positions of the imidazopyrazinone ring.

Although early variants of marine luciferases gained some success, they did not come under the spotlight until the development of the NLuc luciferase in 2012.^7^ NLuc was engineered from the small subunit of a CTZ-utilizing *Oplophorus* luciferase (Oluc)^8^ and particularly optimized for a synthetic CTZ analog, FRZ (**Figure 1**). The photon production rate of the NLuc-FRZ luciferase-luciferin pair is two orders of magnitude higher than RLuc-CTZ and FLuc-D-luciferin.^7^ Moreover, NLuc exhibits a reduced molecular size (19 kDa as opposed to 61 kDa of FLuc and 36kDa of RLuc), high physical stability, high activity in both intracellular and extracellular space, and resistance to domain splitting and circular permutation.^6,7,9,10^ As a result, NLuc is being broadly used in *in vitro* and cellular assays, and a large number of NLuc-based bioluminescent indicators have been developed.^11-18^

Despite the progress, the blue emission (∼450nm) of the NLuc-FRZ pair limits its widespread usage *in vivo*. Short-wavelength light suffers from strong scattering, absorption, and attenuation when it travels through mammalian tissue.^19^ To red-shift the emission of NLuc toward the intravital optical imaging window (600-950 nm), researchers have explored several strategies. First, NLuc has been genetically fused to fluorescent proteins (FPs) with longer-wavelength emission so that a nonradiative <u>b</u>ioluminescence <u>r</u>esonance <u>e</u>nergy transfer (BRET) process occurs to generate red-shift.^16,17,20-22^ The second strategy still relies on BRET, but synthetic small-molecule fluorophores are energy transfer acceptors. In these constructs, NanoLuc or its circularly permuted variants are fused with self-labeling protein tags (e.g., Halo tag), to which synthetic dyes can be covalently tethered.^23-25^ A wide range of bioluminescence colors can thus be obtained by changing the hues of the labeling dyes. However, the dye labeling step adds considerable technical complexity, particularly when applying the technology *in vivo*. In addition, the two strategies mentioned above inevitably increase the molecular size and decrease the folding robustness of the luciferase enzyme. The third approach utilizes quantum dots as BRET acceptors.^26^ NLuc has to be conjugated to quantum dots *in vitro* before being loaded into cells and animals, so the enzyme cannot be used as a genetically encoded label. Finally, the fourth approach is not based on BRET but uses chemically modified CTZ analogs. The functional groups extending through the C2, C6, and C8 positions of the imidazopyrazinone core of CTZ (**Figure 1**) are important for the photophysics of the bioluminescence reaction. Previous studies have reported a series of CTZ analogs showing bioluminescence spanning a broad spectrum in the presence of NLuc.^20,21,24,27^ In particular, extending the substrate conjugation by putting aryl moieties to the C8 position was highly effective for red-shifting bioluminescence.^20,21,24^ An obvious caveat of this approach is that the substrate structural change may impede the host-guest interaction, catalysis, and photon production efficiency, practically resulting in reduced bioluminescence intensity. Our group previously showed that the NLuc luciferase could be engineered to accommodate red-shifted substrates. We have successfully developed teLuc and LumiLuc from NLuc for pairing with CTZ analogs, DTZ and 8pyDTZ, resulting in bright teal (∼500 nm) and chartreuse (∼520 nm) colors, respectively (**Figure 1 & Figure 2a**).^20,21^

**Figure 2.**
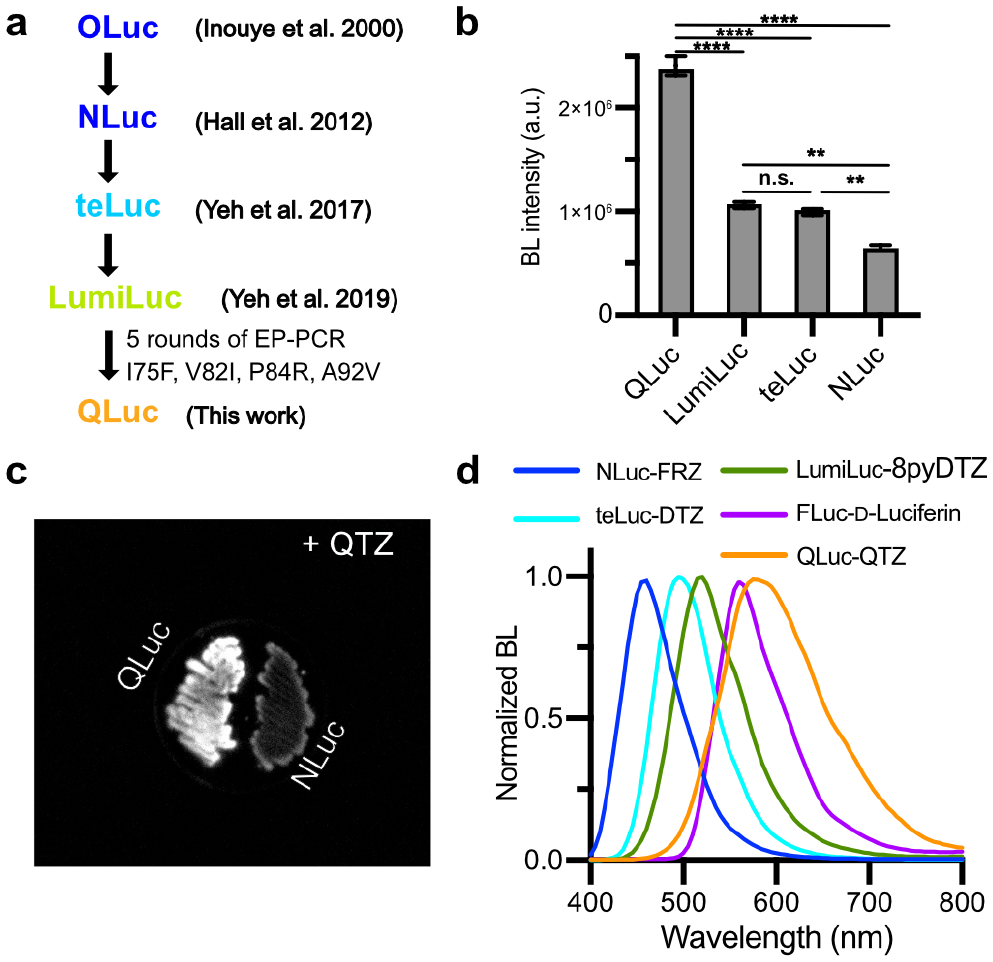
Engineering of the QLuc luciferase. (a) Procedures showing the development of QLuc from OLuc. (b) BL intensity comparisons of QLuc, LumiLuc, teLuc, and NLuc in *E. coli* lysates in the presence of QTZ. Values are expressed as mean ± s.e.m. of three technical replicates. *P* values were determined by one-way ANOVA with Tukey’s multiple comparisons test (\*\*\*\**P*< 0.0001; \*\**P*< 0.01; n.s, not significant, *P*> 0.05). (c) BLI of *E. coli* cells expressing QLuc or NLuc on the LB agar upon spraying a QTZ solution. (d) Normalized BL emission spectra of the indicated luciferase-luciferin pairs.

Antibodies and antibody fragments have superior targeting specificity and have been broadly exploited for antigen recognition. Immunofluorescence imaging with fluorescently labeled antibodies or antibody fragments is a widely employed method for detecting specific antigens in monolayer cell cultures or thin tissue slices.^28^ Meanwhile, immunoPET based on antibodies or antibody fragments labeled with positron (β^+^)-emitting radionuclides is the method of choice for molecular imaging in live animals and humans.^29^

Due to its excellent sensitivity and quantitative capability, ImmunoPET has been used for biomarker visualization, disease diagnosis and staging, imaging-guided therapies, and the preclinical development of new treatments.^30^ However, immunoPET needs expensive and specialized facilities, including PET scanners, radionuclide generators or cyclotrons, and radiolabeling and purification equipment. Furthermore, radiation risks during agent preparation and imaging acquisition further limit the application breadth of immunoPET.

Herein, we report the development of a QTZ luciferin and a paired NLuc variant, namely QLuc, for bright bioluminescence with peak emission at ∼ 585 nm. Leveraging QLuc’s small molecular size, ATP-independency, and red-shifted emission, we created a fusion protein between QLuc and an anti-Her2 scFv and performed BLI of Her2(+) tumors in a xenograft mouse model. We suggest naming this noninvasive molecular imaging method “immunoBLI”, which resembles immunofluorescence imaging and immunoPET but is more suited for animal studies in basic and preclinical research.

## RESULTS

### Design and synthesis of QTZ

Compared to CTZ and FRZ, substrates with extended conjugations through the C8 position of the imidazopyrazinone core may cause red-shifted BL when paired with NLuc.^21,24^ Based on our previous work,^20^ we designed a structural analog of 8pyDTZ by further expanding the conjugation system at the C8. We named this new compound QTZ for its 4-quinolinyl substitution (**Figure 1**). We synthesized QTZ from commercially available compounds by modifying the procedures we previously used to prepare DTZ (**Supporting Information, Scheme S1 & Experimental Methods**).^21^ Briefly, we used a Suzuki cross-coupling for regioselective installation of the 4-quinoline substitution at the C-8 position. Next, we conducted the second Suzuki coupling reaction to conjugate a phenyl group at the C-6 position. Finally, the afforded intermediate was further cyclized with phenyl α-ketoacetal in an acidic condition to yield QTZ.

### Directed evolution of QLuc for enhanced brightness

We preliminarily evaluated the bioluminescence of QTZ with the NLuc enzyme. To our delight, we observed an emission peak centered at ∼585 nm. Thus, compared to FRZ and 8pyDTZ, QTZ caused ∼130 and 65 nm red-shifted emission, respectively. We further examined the bioluminescence of QTZ in the presence of two other NLuc mutants, teLuc and LumiLuc (**Figure 2ab**). Among these three luciferases, LumiLuc gave the highest intensity. The result is not surprising since QTZ is structurally closer to 8pyDTZ than DTZ or FRZ (**Figure 1**). We next performed directed evolution with LumiLuc. We introduced random mutations with error-prone polymerase chain reactions (EP-PCR) and screened the resultant library in *Escherichia coli* colonies for increased brightness using a modified BLI system. (**Figure S1**). After five rounds of directed evolution, we arrived at QLuc with four additional mutations (I75F, V82I, P84R, A92V) from LumiLuc (**Figure S2**). In *E. coli* cell lysates, the brightness of QLuc was ∼ 5-fold of NLuc in the presence of the QTZ substrate (**Figure 2b**). To compare the brightness more intuitively, we expressed NLuc or QLuc in *E. coli* cells streaked on the lysogeny broth (LB) agar in a Petri dish. After a QTZ solution was sprayed over the bacteria, images were acquired in a dark box, and much brighter BL from QLuc was observed (**Figure 2c**).

We further performed assays with purified enzymes and used the NLuc-FRZ pair for comparison (**Figure S3**). The photon production rate of QLuc-QTZ, presented as a relative rate (*k*_cat_) for maximal photon production, was ∼24% of NLuc-FRZ and 2.3-fold of NLuc-QTZ. Under our assay conditions, the Michaelis constant (*K*_M_) values were 1.11, 1.51 and 1.17 µM for NLuc-FRZ, NLuc-QTZ and QLuc-QTZ, respectively (**Table S1**). Thus, in terms of the catalytic efficiency (*k*_cat_/*K*_M_) toward QTZ, QLuc was 3-fold more effective than NLuc.

We next compared the normalized emission spectrum of QLuc-QTZ with several other popular lucif-erase-luciferin pairs (**Figure 2d**). QLuc-QTZ is among the most red-shifted marine luciferase derivatives. It has a broad emission peak, and 52% of its total emission is above 600 nm (**Table S2**). Impressively, it is even more red-shifted than FLuc-D-luciferin, an ATP-dependent insect luciferase reporter system popular for animal imaging. Further *in vitro* assays revealed that the QLuc-QTZ pair was 73-fold brighter than FLuc-D-luciferin at pH 8 (**Figure S4**). The red-shifted emission profile and reasonably high brightness of QLuc-QTZ suggest its promise for *in vivo* imaging.

### Evaluation of the brightness of QLuc-QTZ in cultured mammalian cells

To further test QLuc in mammalian cells, we transiently transfected human embryonic kidney (HEK) 293T cells. HEK 293T cells expressing NLuc, teLuc, or LumiLuc were used for comparison. We added ∼ 10,000 dissociated cells into each well in a 96-well plate and supplemented the corresponding luciferins to a final concentration of 50 μM. To simulate light absorption by mammalian tissue, we imaged the bioluminescence over time through a far-red filter (∼ 600-700 nm). Under this experimental condition, the brightness of QLuc-QTZ outperformed other tested luciferase-luciferin pairs at the beginning and during the duration of the experiment (**Figure 3a**). QLuc-QTZ gave persisting bioluminescence emission, and its integrated far-red signal over the 15-min time window was ∼ 2.1-fold of LumiLuc-8pyDTZ, 2.8-fold of teLuc-DTZ, and 6.4-fold of NLuc-FRZ (**Figure 3bc**). The results suggest that QLuc-QTZ may give bright and long-lasting signals for mammalian cell imaging.

**Figure 3.**
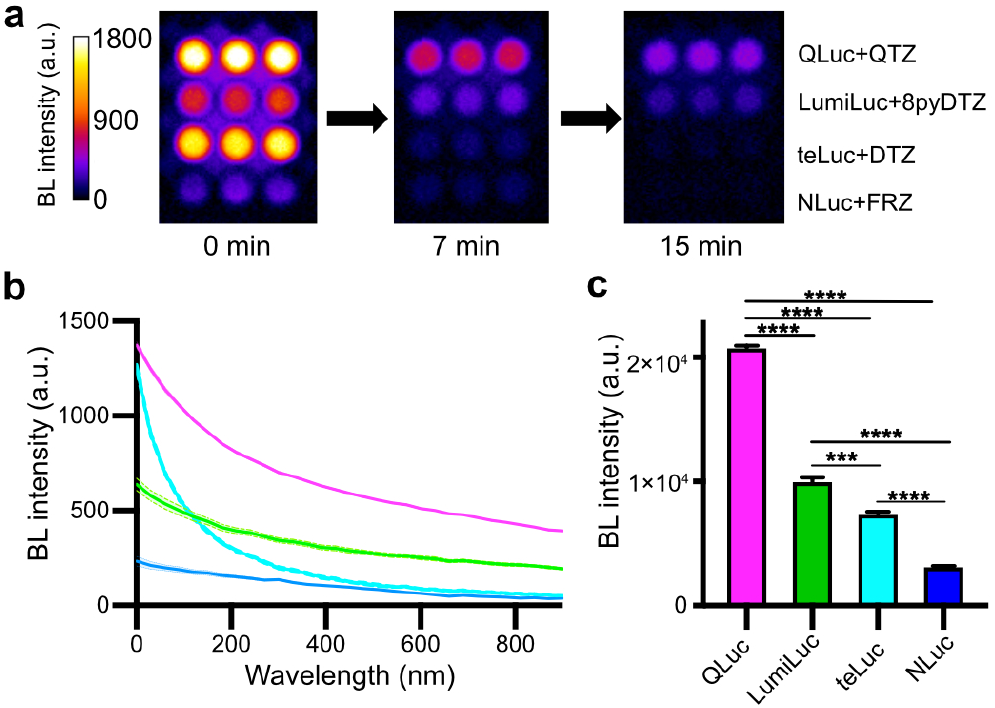
Comparison of the far-red emission of QLuc with several other luciferases in HEK 293T cells. (a) A series of pseudocolored BL photographs of HEK 293T cells in wells of a 96-well plate. The images were acquired with an Andor EMCCD camera through a far-red filter (600-700 nm). (b) The timecourse of the bioluminescence intensities, shown as mean ± s.e.m of three technical replicates. (c) Comparison of the integrated bioluminescence intensity over 15 minutes. Values are expressed as mean ± s.e.m. of three technical replicates. *P* values were determined by one-way ANOVA with Tukey’s multiple comparisons tests (\*\*\*\**P*< 0.0001; \*\*\**P*< 0.001).

### QLuc for imaging deep-tissue targets in vivo

Encouraged by the promising results from the *in vitro* and cellular assays, we next turned to a broadly utilized mouse liver model for evaluating the QLuc-QTZ luciferase-luciferin pair for deep-tissue imaging *in vivo*. We and others previously used a hydrodynamic transfection method to deliver luciferase genes to the liver in mice, but the procedure was highly dependent on operators, resulting in notable experimental variations.^17,21^ We thus revised the process using the systemic injection of adeno-associated viruses (AAVs) containing a liver-specific promoter. Briefly, we first packaged AAVs with the expression of QLuc, NLuc, or teLuc driven by the liver-specific thyroxine-binding globulin (TBG) promoter.^31^ Next, an equal amount of AAVs (i.e., 100 µL at the viral titer of 5×10^12^ gene copies/mL) was administered to mice through the tail vein. The use of the TBG promoter restricted the expression of the luciferases in the transduced liver. We performed the brightness comparison three weeks post-viral administration by administering an equal amount (0.1 µmol) of the substrates intraperitoneally (**Figure 4a**). Mice were imaged from the dorsal side, with thick tissue above the liver. The initial bioluminescence of the QLuc-QTZ pair was ∼ 50% of teLuc-DTZ and ∼ 1.5-fold of NLuc-FRZ (**Figure 4bc**). The intensity of QLuc gradually increased and reached a plateau ∼ 10 min post the substrate injection. Impressively, the high intensity of QLuc was maintained for more than 30 min. In contrast, teLuc and NLuc signals faded quickly. Compared to either teLuc-DTZ or NLuc-FRZ, the signal from QLuc-QTZ was much higher at the 25^th^ and 50^th^ min post substrate injections, respectively (**Figure 4d**). As for the bioluminescence signals integrated over the 50-min experimental period, QLuc was ∼2.5- and 2-fold brighter than NLuc and teLuc, respectively (**Figure 4e**). Collectively, these data indicate that the QLuc-QTZ pair has excellent deep-tissue performance and is suited for *in vivo* BLI in small mammals.

**Figure 4.**
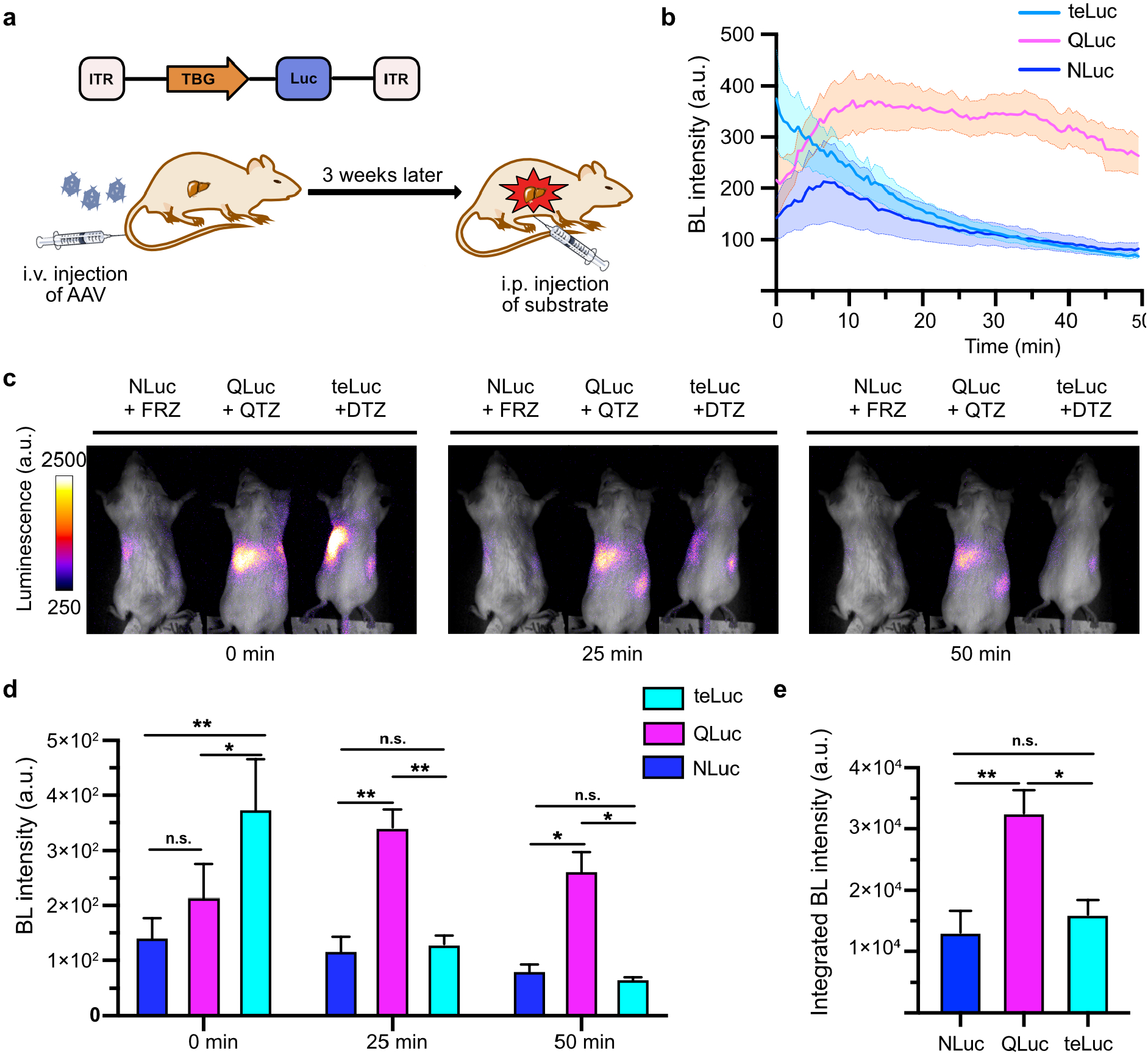
Comparison of QLuc with NLuc and teLuc for BLI of the liver in mice. (a) Schematic diagram of the genetic constructs for the luciferase genes driven by the liver-specific thyroxine-binding globulin (TBG) promoter, and the experimental timeline for viral transduction and imaging. (b) Timecourse of BL intensities over the liver regions. Values are expressed as mean ± s.e.m. of five biological replicates in each group. (c) A series of pseudocolored BL images, showing the high brightness and slow decay of QLuc-QTZ. (d) Comparison of BL intensities at the indicated time points. Values are expressed as mean ± s.e.m. of five biological replicates. *P* values were determined by two-way ANOVA with Tukey’s multiple comparisons test (\*\**P*< 0.01; \**P*< 0.05; n.s, not significant, *P*> 0.05). (e) Comparison of bioluminescence intensities integrated over the 50-min period. Values are expressed as mean ± s.e.m. of five biological replicates. *P* values were determined by one-way ANOVA with Tukey’s multiple comparisons test (\*\**P*< 0.01; \**P*< 0.05; n.s, not significant, *P*> 0.05).

### QLuc as a nonradiative tracer for antibody-based molecular imaging in vivo

The activity of QLuc is independent of ATP, and thus, it remains functional in the extracellular space where ATP concentrations are low.^6^ Because of its bright and red-shifted emission and excellent photon penetration in mammalian tissue, we envisaged that QLuc could be used to tag antibodies or antibody fragments for imaging specific cell-surface markers *in vivo*. To test this hypothesis, we genetically fused QLuc to an anti-Her2 scFv derived from the trastuzumab (Herceptin) antibody (**Figure 5a, Table S3 & Table S4**). The variable domains of trastuzumab (Herceptin) light and heavy chains (V_L_ and V_H_) were linked with four copies of a GGGGS linker, and the C-terminus of the V_H_ was fused to the N-terminus of QLuc through a previously reported protease-resistant floppy linker.^32^ A His_6_ tag was further appended to the C-terminus of QLuc for affinity purification. We expressed and purified the fusion protein from *E. coli*. A SHuffle T7 bacterial strain, engineered from the *E. coli* B strain with glutaredoxin and thioredoxin reductase knockout and disulfide bond isomerase overexpression, was used for protein expression, since we expected it to promote the formation of the disulfide bonds in the scFv. Indeed, the fusion protein was soluble in the cytoplasm, and we gained ∼ 30 mg of the protein from a liter of liquid culture without optimizing the expression and purification conditions.

**Figure 5.**
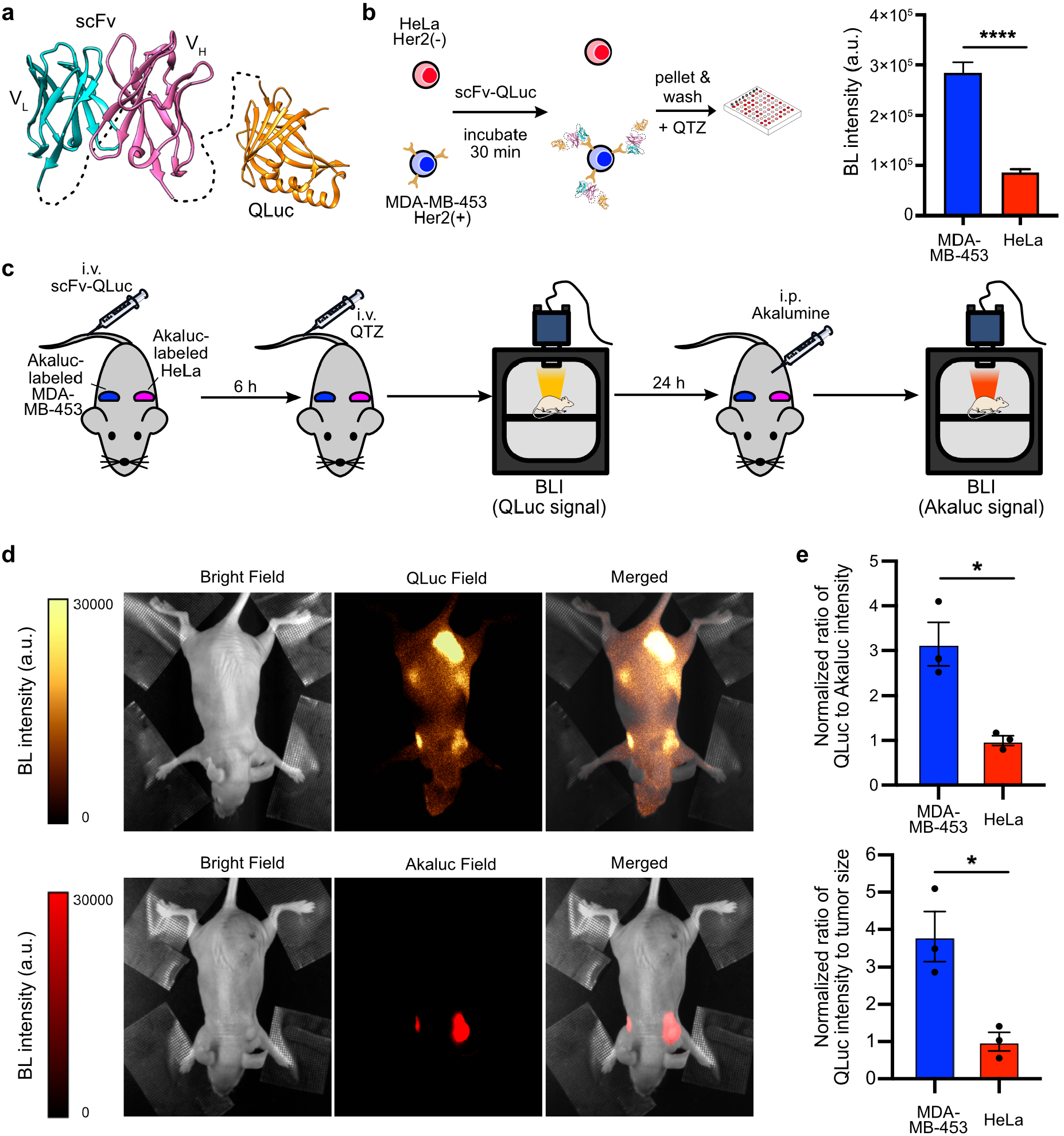
ImmunoBLI of the tumor-associated Her2 antigen using an scFv-QLuc fusion protein. (a) Domain arrangement of the anti-Her2 scFv-QLuc fusion construct. An scFv derived from trastuzumab (Herceptin) was genetically fused to the N-terminus of QLuc, and the fusion construct was expressed and purified from *E. coli*. The illustration was created based on PDB entries 1N8Z and 7MJB. (b) Workflow to test the enrichment of the anti-Her2 scFv-QLuc fusion protein at the surface of cultured Her2(+) MDA-MB-453 cells using a BL immunoassay. The plate reader readings of the BL intensities are presented as mean ± s.e.m. of three technical replicates. *P* values were determined by unpaired t-test (\*\*\*\**P*< 0.0001). (c) Workflow to use the fusion protein for *in vivo* ImmunoBLI. The Akaluc-Akalumine luciferase-luciferin pair, biochemically orthogonal to QLuc-QTZ, was used as a stable intracellular label of Her2(+) and Her2(-) cells. (d) Representative pseudocolored BL images of mice for QLuc-QTZ signals (top) and Aka-luc-Akalumine signals (bottom), showing the biodistribution of the anti-Her2 scFv-QLuc fusion protein in relation to the Akaluc-labeled tumors. (e) Quantitative analysis of QLuc signals over the regions of the Her2(+) and Her2(-) tumors, showing the specific accumulation of the scFv-QLuc fusion protein in Her2(+) tumors. Values are normalized to Akaluc intensities (top) or tumor sizes measured with a caliber (bottom) and expressed as mean ± s.e.m. of three biological replicates. *P* values were determined by unpaired t-test (\**P*< 0.05).

To prove that the prepared scFv-QLuc fusion protein is functional, we incubated the proteins with equal numbers of Her2(+) MDA-MB-453 cells and Her2(-) HeLa cells. After washing, emission intensities were quantified with a plate reader after supplementing the QTZ substrate (**Figure 5b**). The signal from MDA-MB-453 cells was ∼ 3.3-fold higher than HeLa cells. Thus, the bioluminescence immunoassay suggests that the scFv-QLuc fusion protein was explicitly enriched at the surface of the Her2(+) cells.

We next used the scFv-QLuc fusion protein to image Her2(+) tumors in a xenograft mouse model. To facilitate the tracking of implanted cancer cells, we first modified MDA-MB-453 and HeLa cell lines to stably express an Akaluc luciferase. Akaluc was recently engineered from FLuc, with peak emission around 650 nm in the presence of a synthetic Akalumine luciferin.^33^ Akaluc was reported for superior *in vivo* sensitivity. Because it is an ATP-dependent luciferase, we expressed Akaluc in the cytosol of the cancer cells. Three weeks after tumor inoculation in the left and right shoulders in NU/J nude mice with the Akaluc-labeled cells, we performed BLI (**Figure 5c**). We injected 1 mg of the scFv-QLuc fusion protein into each mouse via the tail vein. Next, we waited for 6 h to allow protein circulation, enrichment, and partial clearance, before performing BLI with anesthetized mice right after an intravenous (i.v.) administration of QTZ. On the next day, when luminescence signals from QLuc and QTZ were entirely dissipated, we intraperitoneally (i.p.) administered Akalumine to mice and acquired another set of images for signals from Akaluc. The first set of images gave information on the biodistribution of the scFv-QLuc fusion protein, whereas the second set of images highlighted the locations of tumors. To our delight, we observed strong QLuc emission from where MDA-MB-453 cells were inoculated (**Figure 5d**). Compared to the intensity of residual QLuc in normal tissues, a signal-to-background ratio (S/B) of >4 was obtained. We also observed strong QLuc emission from the low abdomen around the kidney and bladder regions, corroborating previous immunoPET studies showing the excretion of scFvs via the kidney into the urine^34,35^ and our work showing the high activity of NLuc-derived luciferases in blood and urine.^6^ There was a low level of scFv-QLuc enrichment at the location where HeLa cells were inoculated. This nonspecific enrichment is known as the enhanced permeability and retention (EPR) effect of tumors.^36^ We further quantitatively compared the ability of the two types of tumors for enriching the scFv-QLuc fusion protein. Because the MDA-MB-453 and HeLa tumors showed notably different volumes, we normalized the QLuc signals from the tumors with either the Akaluc BL intensities or tumor sizes measured with a caliber (**Figure 5e**). The Her2(+) MDA-MB-453 tumors were over 3-fold more effective in enriching the scFv-QLuc fusion protein. Collectively, the experiments demonstrate an “immunoBLI” approach, which uses QLuc as a nonradiative tracer for antibody-based molecular imaging of tumor-associated antigens in a mouse model. Furthermore, the results presented here support the feasibility of using QLuc and Akaluc for orthogonal two-population BLI in vivo.

## DISCUSSION AND CONCLUSIONS

In this work, we designed and chemically synthesized QTZ, a CTZ analog with the 4-quinolinyl substitution at the C8 position of the imidazopyrazinone core. We further engineered a QLuc luciferase from LumiLuc to pair with QTZ. The resultant QLuc-QTZ luciferase-luciferin pair showed a peak emission at ∼585 nm, and more than half of its total emission was above 600 nm. The intrinsic brightness of QLuc-QTZ was about a quarter of NLuc-FRZ but still over 70-fold higher than FLuc-D-luciferin. Because of its red-shifted emission profile and reasonably high brightness, QLuc-QTZ significantly improved tissue penetration in a mouse model with luciferases expressed in the liver. More impressively, the BL of QLuc sustained much longer than other tested marine luciferases, making QLuc an exceptionally promising reporter for *in vivo* BLI in small mammals. We expect further studies to continue improving this lucifer-ase-luciferin pair (*e*.*g*., increase substrate solubility and redder and brighter emission)

Animal models are indispensable for understanding new mechanisms in physiological and pathological processes, as well as testing new therapies for diseases. Antibody-based molecular imaging in mammalian model organisms urgently needs the development of inexpensive and convenient nonradiative tracers. Luciferases, which may be used to label antibodies or antibody fragments, have the potential to fill the niche. However, ATP-dependent insect luciferases only work under conditions with plenty of ATP (e.g., intracellular space). On the other hand, marine luciferases are enzymatically active in the extracellular space, but conventional marine luciferases, such as RLuc, have blue emission, low brightness, and limited serum stability, so the luciferase-antibody fusion proteins do not give reliable sensitivity *in vivo*.^37^ The photon production rate of the recently developed NLuc-FRZ luciferase-luciferin pair is much improved from RLuc-CTZ. Also, NLuc and its mutant luciferases are small and stable, and they retain high activity in complex biological media, such as serum and urine. Leveraging the favorable features of QLuc inherited from NLuc and the drastically red-shifted emission of QLuc, we used QLuc as a nonradiative tracer for labeling antibody fragments and visualizing tumor-associated antigens in a xenograft mouse model. This immunoBLI method is attractively positioned between immunofluorescence imaging and immunoPET. It is a convenient and inexpensive molecular imaging tool for animal models in basic and preclinical research.

This study used an scFv for target recognition because the fast penetration of the scFv to tumors and its fast clearance from circulation is an advantage for imaging applications.^38^ We developed a scalable and straightforward method to express the scFv-QLuc fusion protein in the *E. coli* cytosol with a high yield. Periplasmic expression or more complex expression hosts are not needed. We thus believe other researchers can broadly adapt our approach.

NLuc has been successfully used to label full-length antibodies for *in vitro* BL assays.^39^ Therefore, it may be possible to follow a similar procedure to generate QLuc-labeled full-length antibodies. Since full-length antibodies have prolonged pharmacokinetics,^38^ such fusion proteins will be less suited as *in vivo* imaging agents. However, the fusion proteins may be valuable tools for studying the pharmacokinetics and therapeutic effects of antibodies in animal models, accelerating the preclinical development of antibody-based drugs.

The applications of this amber-light emitting QLuc may be further explored in many other scenarios. For example, its small molecular size may facilitate the integration of the luciferase gene into a viral genome for investigating viral replication and propagation *in vivo*. Moreover, the photophysical properties of QLuc make it a desirable start-point for developing point-of-care assays since the red-shifted emission is less absorbed by hemoglobin in the blood. Furthermore, QLuc may pave the way for developing red-shifted luminescent indicators, facilitating the noninvasive visualization of biochemical events in deep sites in animals. In addition, QLuc may be a suitable BRET donor for far-red and near-infrared (NIR) fluorophores. Thus, it may be used to study molecular interactions or develop new BRET-based luminescent indicators. Finally, can QLuc or QLuc-based immunoBLI be applied to humans? We can imagine potential clinical applications, but the toxicity of the substrate and the immunogenicity of the luciferase in humans must be carefully investigated.

## ASSOCIATED CONTENT

### Supporting Information

Figures S1-S4, including experimental procedure illustration, sequence alignment, enzyme assay results, and additional BL spectra; Tables S1-S4 for luciferase photophysical properties and DNA, protein and oligonucleotide sequences; Scheme S1; experimental methods; and NMR spectra for newly synthesized compounds.

## AUTHOR INFORMATION

### Notes

HA and YX are listed as inventors of a patent or a patent application covering some luciferase and luciferin variants described in this work.

### Author Contributions

HA conceived the project. YX synthesized compounds, engineered QLuc with the help of MSR, performed characterization, and prepared AAVs and scFv-QLuc. YZ, XL, and YX performed animal experiments. ZL established Akaluc-labeled stable cell lines. XT provided FRZ for comparison experiments. YX analyzed data and prepared figures. HA and YX wrote the manuscript with inputs from other authors.

## ACKNOWLEDGMENTS

We thank Hao Zhang and Tianchen Wu for their technical assistance, and Drs. Wladek Minor and Ivan Shabalin for providing the crystal structure of NLuc-R164Q. Research reported in this publication was supported in part by the University of Virginia Start-up Fund and National Institutes of Health grants (R01DK122253, RF1AG077773, and R01GM129291) to HA.

