## Supporting Information for "An Engineered Amber-Emitting Nano Luciferase and Its Use for Immunobioluminescence Imaging in Vivo"

**This PDF file includes:**

Figures S1 to S4
Tables S1 and S4
Scheme S1
Experimental Methods
NMR Spectra

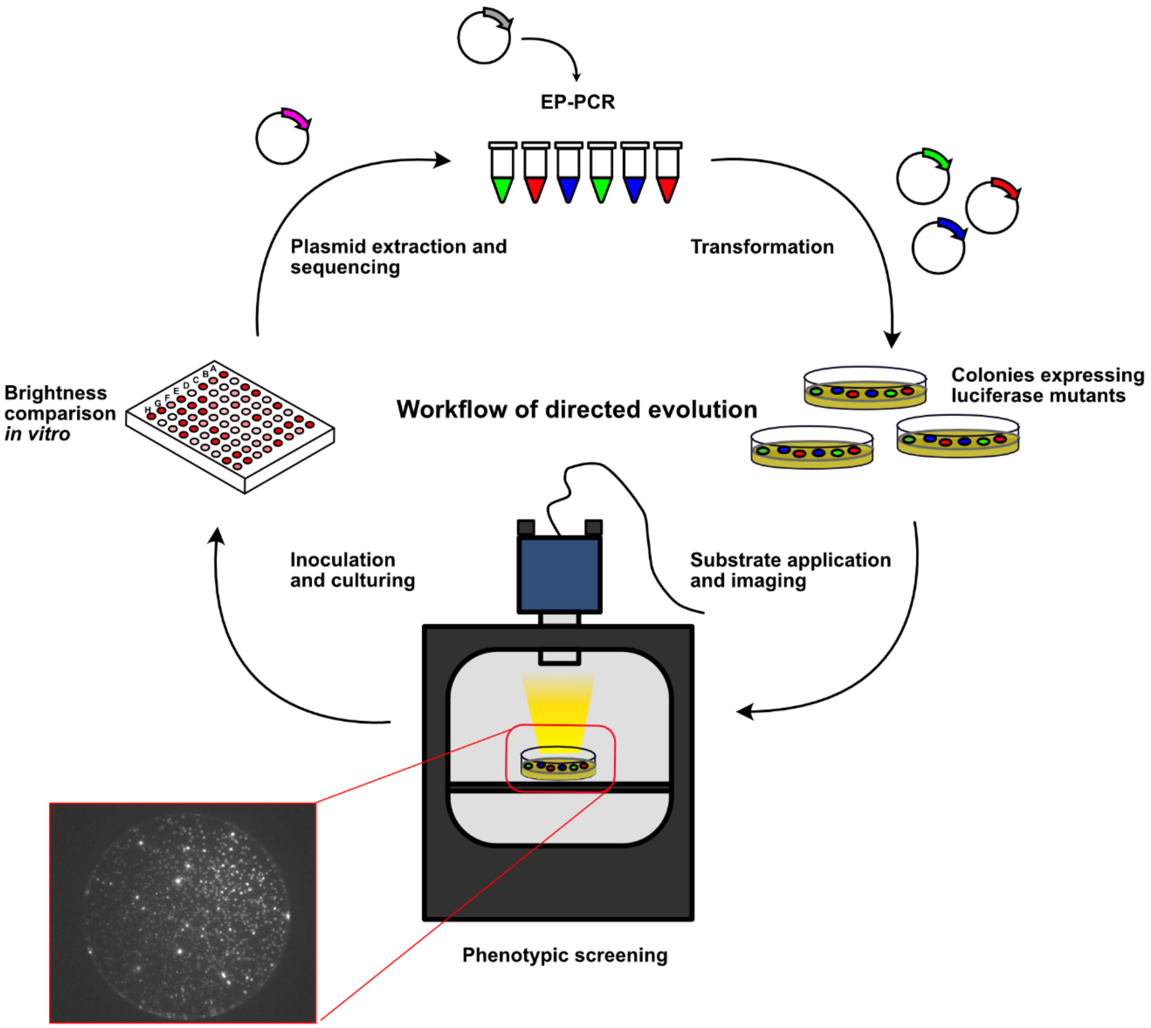


**Figure S1. Illustration of the directed evolution process to derive QLuc.** Random mutations were introduced with error-prone PCR (EP-PCR). *E. coli* libraries expressing luciferase mutants were subjected to phenotypic screening upon spraying the QTZ substrate to colonies in Petri dishes. BL images were acquired using a cooled CCD camera in a dark box. Bright colonies were selected, cultured, lysed, and verified in 96-well plates. The brightest mutants were used for subsequent rounds of directed evolution.

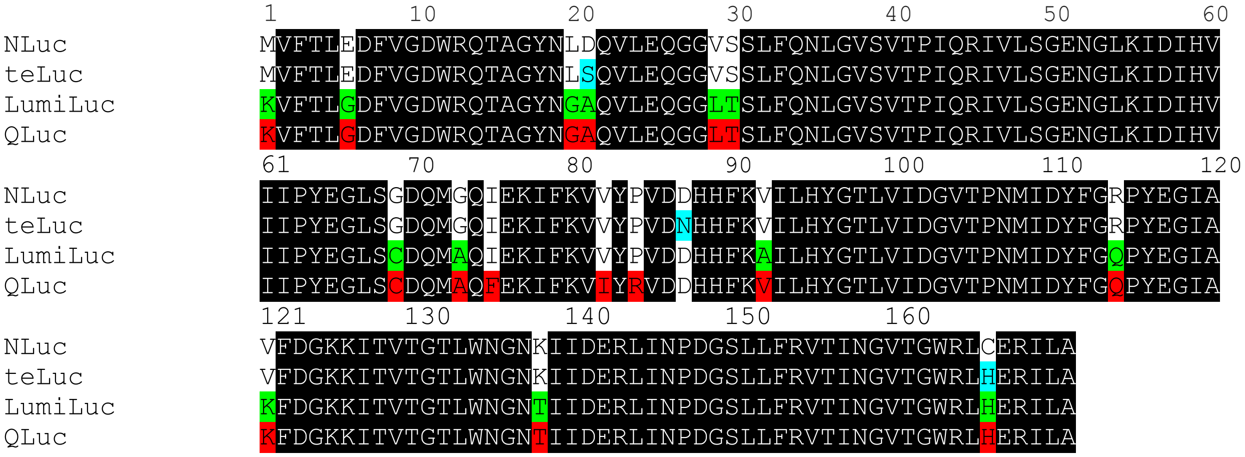


**Figure S2. Sequence alignments of NLuc, teLuc, LumiLuc and QLuc.** Mutations in teLuc, LumiLuc, and QLuc compared to NLuc are highlighted in teal, green, and red, respectively. Residues in this figure are numbered according to PDB ID 7MJB.

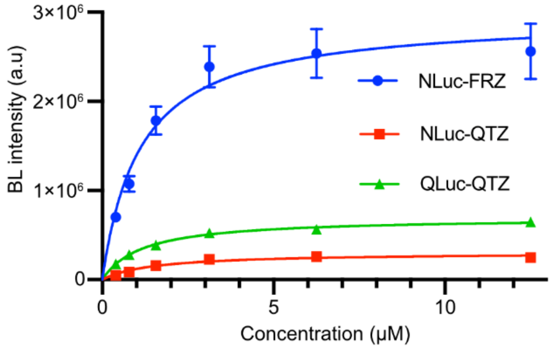


**Figure S3. Enzyme assays to determine the relative photon production rates ($k_{cat}$) and apparent Michaelis constants ($K_M$) of QTZ in the presence of the indicated luciferases.** As the same concentrations of purified luciferases were used, the relative $k_{cat}$ values were derived from the maximal BL intensity values. Mean values are presented with error bars representing s.e.m. (n=3).

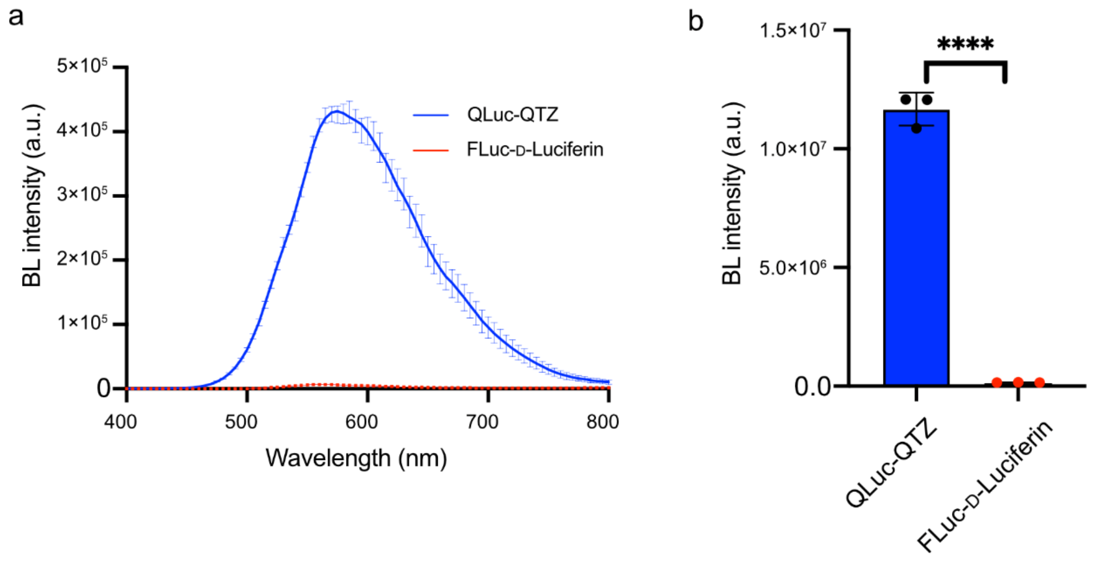


**Figure S4. Comparisons of QLuc-QTZ and FLuc-D-Luciferin using purified enzymes.** (a) Emission spectra measured from 400 nm to 800 nm under the following condition: 100 nM luciferase, 150 mM NaCl, 1.5 mM ATP, 5 mM $MgSO_4$, 30 mM Tris-HCl, pH 8, 100 µM D-luciferin or 20 µM QTZ. (b) Comparison of the brightness (calculated as the integration of instrumental counts over wavelengths from 400 to 800nm). Mean values are presented with error bars representing s.d. (n=3). *P* values were determined by unpaired t-test (*****P*< 0.0001).

**Table S1. The relative photon production rates ($k_{cat}$) and apparent Michaelis constants ($K_M$) of the indicated luciferase-luciferin pairs**.

| | NLuc-FRZ | NLuc-QTZ | QLuc-QTZ |
|---|---|---|---|
| Relative $k_{cat}$ | 1 | 0.103 | 0.236 |
| $K_M$ (μM) | 1.11 | 1.51 | 1.17 |
| Relative $k_{cat}/K_M$ | 1 | 0.076 | 0.224 |

**Table S2. Emission wavelengths of luciferase-luciferin pairs described in this work.** The percentage of photons over 600 nm within the total emission is also presented since photons over 600 nm are generally regarded to have better tissue penetration.

| Luciferase | Luciferin | Emission peak | Photons > 600 nm |
|---|---|---|---|
| NLuc | FRZ | 455 nm | 2% |
| teLuc | DTZ | 500 nm | 4% |
| LumiLuc | 8pyDTZ | 520 nm | 11% |
| QLuc | QTZ | 585 nm | 52% |
| FLuc | D-Luciferin | 560 nm | 35% |

**Table S3. DNA and protein sequences of the QLuc luciferase and the anti-Her2 scFv-QLuc fusion construct.** For the latter construct, the $V_L$ and $V_H$ chains of trastuzumab are highlighted in yellow and green, respectively. The floppy linkers are highlighted in grey. The QLuc portion is highlighted in teal.

| | |
|---|---|
| DNA sequence of QLuc | aaagtcttcactctcggggattttgttggggactggcgacagacagccggctacaaccaggctcaagtccttgaacagggaggtttgacc agtttgtttcagaacctcggggtgtccgtaactccaatccaaaggattgtcctgagcggtgaaaatgggctgaagatcgatatccatgtcatc atcccgtatgaaggtctgagctgcgaccaaatggcccagttcgaaaaaatttttaaggtgatataccgtgtggatgatcatcactttaaggtg atcctgcactatggcacactggtaatcgacggggttacgccgaacatgatcgactatttcggacagccgtatgaaggcatcgccaagttcg acggcaaaaagatcacagtaacagggaccctgtggaacggcaacacaattatcgacgagcgcctgatcaaccccgacggctccctgct gttccgagtaaccattaacggagtgaccggctggcgtctgcatgaacgtattctggcgtaa |
| Protein sequence of QLuc | KVFTLGDFVGDWRQTAGYNQAQVLEQGGLTSLFQNLGVSVTPIQRIVLSGENGLKIDI HVIIPYEGLSCDQMAQFEKIFKVIYRVDDHHFKVILHYGTLVIDGVTPNMIDYFGQPYE GIAKFDGKKITVTGTLWNGNTIIDERLINPDGSLLFRVTINGVTGWRLHERILA- |
| DNA sequence of anti-Her2 scFv-QLuc fusion antibody | atgggcagcgatattcagatgacccagagcccgagcagcctgagcgcgagcgtgggcgatcgcgtgaccattacctgccgcgcgagc caggatgtgaacaccgcggtggcgtggtatcagcagaaaccgggcaaagcgccgaaactgctgatttatagcgcgagctttctgtatag cggcgtgccgagccgctttagcggcagccgcagcggcaccgattttaccctgaccattagcagcctgcagccggaagattttgcgacct attattgccagcagcattataccaccccgccgacctttggccagggcaccaaagtggaaattaaacgcaccggcggtggcggtagcggt ggcggtggcagcggtggtggcggcagcggcggcggtggtagcgaagtgcagctggtggaaagcggcggcggcctggtgcagccg ggcggcagcctgcgcctgagctgcgcggcgagcggctttaacattaaagatacctatattcattgggtgcgccaggcgccgggcaaag gcctggaatgggtggcgcgcatttatccgaccaacggctatacccgctatgcggatagcgtgaaaggccgctttaccattagcgcggata ccagcaaaaacaccgcgtatctgcagatgaacagcctgcgcgcggaagataccgcggtgtattattgcagccgctggggcggcgatgg cttttatgcgatggattattggggccagggcaccctggtgaccgtgagcagcggcagcaccagcggcagcggcaaaagcagcgaagg caaaggcaaagtcttcactctcggggattttgttggggactggcgacagacagccggctacaaccaggctcaagtccttgaacagggag gtttgaccagtttgtttcagaacctcggggtgtccgtaactccaatccaaaggattgtcctgagcggtgaaaatgggctgaagatcgatatc catgtcatcatcccgtatgaaggtctgagctgcgaccaaatggcccagttcgaaaaaatttttaaggtgatataccgtgtggatgatcatcac tttaaggtgatcctgcactatggcacactggtaatcgacggggttacgccgaacatgatcgactatttcggacagccgtatgaaggcatcg ccaagttcgacggcaaaaagatcacagtaacagggaccctgtggaacggcaacacaattatcgacgagcgcctgatcaaccccgacgg ctccctgctgttccgagtaaccattaacggagtgaccggctggcgtctgcatgaacgtattctggcgctcgagcaccaccaccaccacca ctga |
| Protein sequence of anti-Her2 scFv-QLuc fusion antibody | MGSDIQMTQSPSSLSASVGDRVTITCRASQDVNTAVAWYQQKPGKAPKLLIYSASFLY SGVPSRFSGSRSGTDFTLTISSLQPEDFATYYCQQHYTTPPTFGQGTKVEIKRTGGGGSG GGGSGGGGSGGGGSEVQLVESGGGLVQPGGSLRLSCAASGFNIKDTYIHWVRQAPGK GLEWVARIYPTNGYTRYADSVKGRFTISADTSKNTAYLQMNSLRAEDTAVYYCSRWG GDGFYAMDYWGQGTLVTVSSGSTSGSGKSSEGKGKVFTLGDFVGDWRQTAGYNQAQ VLEQGGLTSLFQNLGVSVTPIQRIVLSGENGLKIDIHVIIPYEGLSCDQMAQFEKIFKVIY RVDDHHFKVILHYGTLVIDGVTPNMIDYFGQPYEGIAKFDGKKITVTGTLWNGNTIIDE RLINPDGSLLFRVTINGVTGWRLHERILALEHHHHHH- |

**Table S4. Oligonucleotides used in this study.**

| Oligo Name | Nucleotide Sequence (5′->3′) |
|---|---|
| pBAD_F_QLuc | gtggacagcaaatgggtcgggatctgtacgacgatgacgataaggatccgagctcgagc |
| pBAD_R_QLuc | gttctgatttaatctgtatcaggctgaaaatcttctctcatccgccaaaacagccaagc |
| pcDNA3_F_QLuc | cgactcactatagggagacccaagcttgccaccatgaaagtcttcactctcggggattttg |
| pcDNA3_R_QLuc | cactatagaatagggccctctagatgcatgctcgagttacgccagaatgcgttcatgc |
| pAAV_F_NLuc | tgtccaggcggccgcgccaccatggtcttcacactcgaagatttcg |
| pAAV_R_NLuc | tcacaaattttgtaatccagaggttgattggatccttacgccagaatgcgttcgc |
| pAAV_R_teLuc | taatccagaggttgattggatccttacgccagaatgcgttcatgcag |
| pAAV_F _QLuc | gtgtccaggcggccgcgccaccatgaaagtcttcactctcgggg |
| pAAV_R_QLuc | tcacaaattttgtaatccagaggttgattggatccgttacgccagaatacgttcatg |
| pET_F1_scFvQLuc | gtttaactttaagaaggagatataccatggatgggcagcgatattcagatgac |
| pET_R1_scFvQLuc | cgctgcttttgccgctgccgctggtgctgccgctgctcacggtcaccag |
| pET_F2_scFvQLuc | gcaccagcggcagcggcaaaagcagcgaaggcaaaggcaaagtcttcactctcgggg |
| pET_R2_scFvQLuc | agtggtggtggtggtggtgctcgagcgccagaatacg |
| Plx208-F-Akluc | tctctggctaactgtcgggatccgccaccatggaagatgccaa |
| Plx208-R-Akluc | ggttgattgtcgacttaacgcgtttacacggcgatcttgcc |

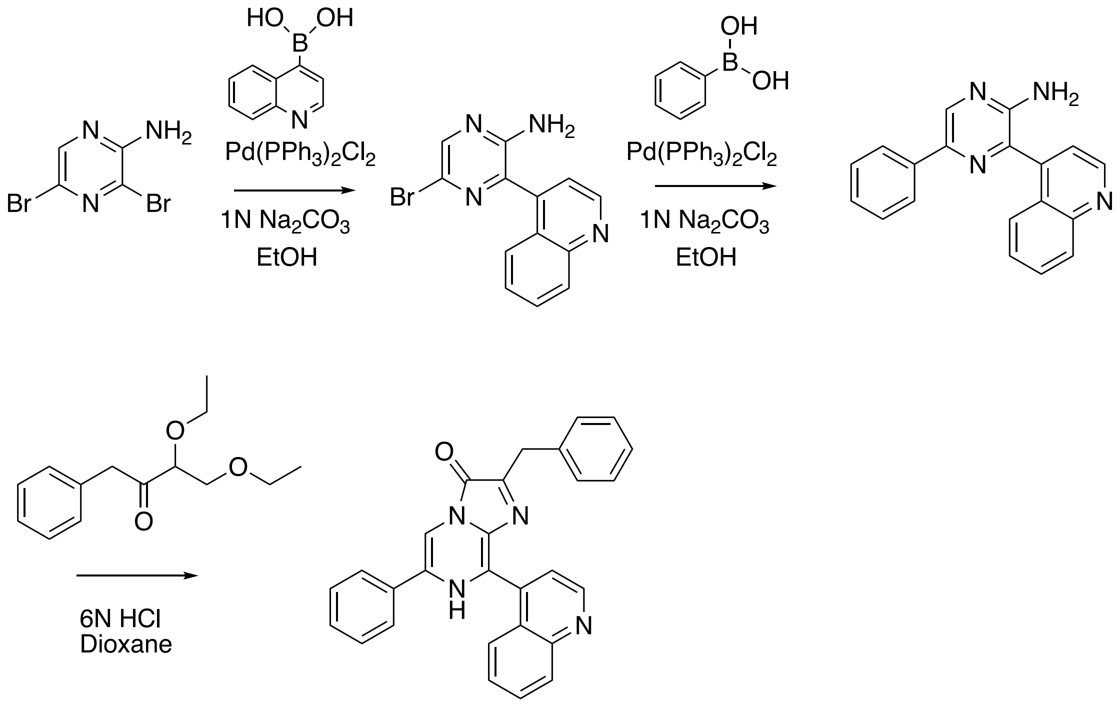


**Scheme S1. Synthetic scheme to derive QTZ.** The tandem Suzuki coupling reactions installed 4-quinolinyl and phenyl groups at the C8 and C6 positions of the imidazopyrazinone core, respectively. The final cyclization process was completed under a strong acid condition to yield QTZ. The total yield for the three steps was ~ 10%.

**Experimental Methods**

**1. Chemical Synthesis**

**1.1 Materials and General Methods**

Unless otherwise noted, all chemicals were purchased from Sigma-Aldrich, Fisher Scientific, or VWR and used without further purification. All $^{1}H$ and $^{13}C$ NMR spectra were collected on a Bruker Avance DRX 600 NMR Spectrometer at the UVA Biomolecular Magnetic Resonance Facility. Chemical shifts (δ) are given with the internal standards: $^{1}H$ (2.50 ppm) and $^{13}C$ (40.0 ppm) for DMSO-d6; $^{1}H$ (3.30 ppm) and $^{13}C$ (49.15 ppm) for methanol-d4. Splitting patterns are reported as s (singlet), bs (broad singlet), d (doublet), t (triplet), dd (doublet of doublets), and m (multiplet). Coupling constants (J) are reported in Hz. Preparative reverse-phase HPLC purifications were conducted on a Waters Prep 150/SQ Detector 2 LC-MS Purification equipped with an XBridge BEH Amide/Phenyl OBD Prep Column (130Å, 5 μm, 30 mm × 150 mm). Lyophilization was performed on a 12-port Labconco freeze dryer equipped with an Edwards RV3 vacuum pump.

**1.2 Synthesis of QTZ**

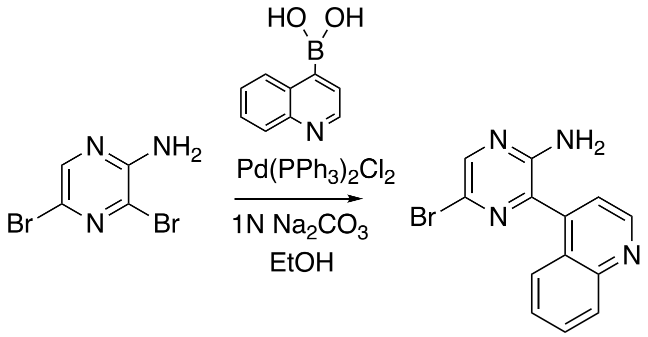


**Compound 1**

To $Pd(PPh_3)_2Cl_2$ (140 mg, 0.2 mmol, 0.1 equiv.) dissolved in 50 mL EtOH, 2-amino-3,5-dibromopyrazine (505 mg, 2 mmol, 1 equiv.),1 N $Na_2CO_3$ solution (4 mL, 4 mmol, 2 equiv.) and quinoline-4-boronic acid (346 mg, 2 mmol, 1 equiv.) were added. The resultant mixture was stirred at 80 °C under argon for 12 h. The solvent was removed in vacuo and the residue was extracted three times with 200 mL EA. The organic layers were pooled and dried over anhydrous $Na_2SO_4$ and then filtered and concentrated in vacuo. The afforded residue was purified by silica column chromatography with elution (EA:HEX= 50:50 ~ 100% EA) to yield Compound 1 as yellowish powder (240 mg, 0.8 mmol, 40%). $^1$H NMR (600 MHz, DMSO-d6) δ 9.01 (d, J = 4.4 Hz, 1H), 8.27 (s, 1H), 8.13 (d, J = 8.5 Hz, 1H), 7.82 (t, J = 7.6 Hz, 1H), 7.65 (d, J = 8.5 Hz, 1H), 7.61 (q, J = 6.7, 5.9 Hz, 3H), 7.55 (d, J = 7.7 Hz, 1H), 6.44 (s, 2H). $^{13}$C NMR (151 MHz, DMSO-*d*$_6$) δ 153.69, 150.86, 148.56, 144.72, 141.67, 136.88, 130.18, 129.91, 129.24, 127.62, 125.91, 125.56, 122.23. ESI-MS ($C_{13}H_{10}BrN_4$): $[M+H]^+$ calcd: 301.00, found: 301.05.

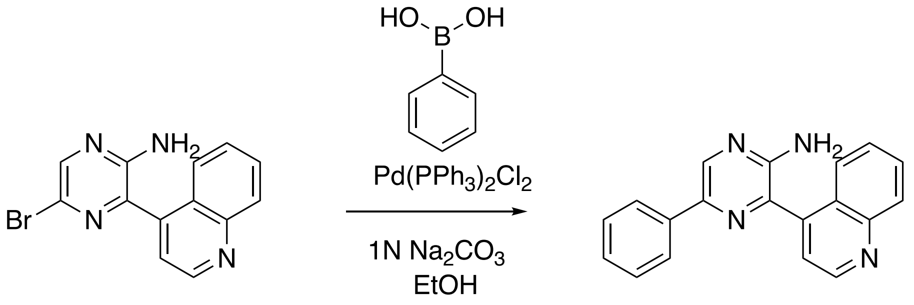


**Compound 1** **Compound 2**

To a solution of $Pd(PPh_3)_2Cl_2$ (56 mg, 0.08 mmol, 0.1 equiv.) dissolved in 20 mL EtOH, 1 N $Na_2CO_3$ solution (1.6 mL, 1.6 mmol, 2 equiv.), phenyl boronic acid (195 mg, 1.6 mmol, 2 equiv.) and Compound 1 (240 mg, 0.8 mmol, 1 eq) were added sequentially. The resultant mixture was stirred at 80 °C under argon for 12 h. The solvent was removed in vacuo and the residue was extracted three times with 200 mL EA. The organic layers were pooled and dried over anhydrous $Na_2SO_4$ and then filtered and concentrated in vacuo. The afforded residue was further purified by silica column chromatography with elution (EA:HEX= 50:50 ~ 100% EA) to yield Compound 2 as yellowish powder (191 mg, 0.64mmol, 80%). $^1$H NMR (600 MHz, DMSO-*d*$_6$) δ 9.01 (d, *J* = 4.3 Hz, 1H), 8.72 (s, 1H), 8.13 (d, *J* = 8.5 Hz, 1H), 7.93 (d, *J* = 7.7 Hz, 2H), 7.79 (t, *J* = 7.6 Hz, 1H), 7.72 (d, *J* = 8.4 Hz, 1H), 7.64 (d, *J* = 4.2 Hz, 1H), 7.60 – 7.51 (m, 1H), 7.40 (t, *J* = 7.6 Hz, 2H), 7.30 (t, *J* = 7.3 Hz, 1H), 6.32 (s, 2H). $^{13}$C NMR (151 MHz, DMSO-*d*$_6$) δ 153.26, 150.97, 148.89, 143.08, 139.91, 139.77, 137.19, 135.91, 130.05, 129.94, 129.18, 128.14, 127.37, 126.42, 125.82, 125.42, 122.22. ESI-MS ($C_{19}H_{15}N_4$): $[M+H]^+$ calcd: 300.13, found: 300.25.

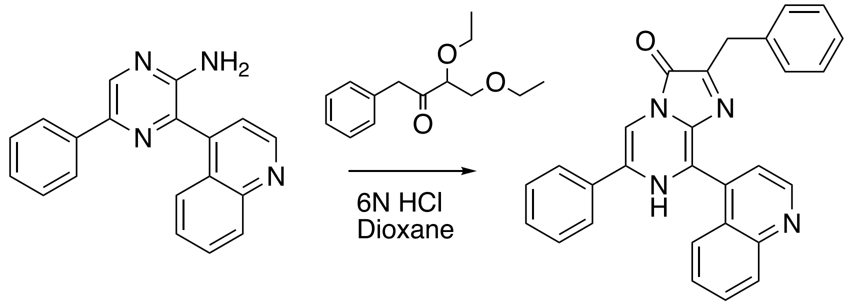

**Compound 2** **QTZ**

To a solution of Compound 2 (30 mg, 0.1 mmol, 1 equiv.) and 1,1-diethoxy-3-phenylpropan-2-one (89 mg, 0.4 mmol, 4 equiv.) in 5 mL degassed 1,4-dioxane, 0.5 mL 6 N HCl (30 equiv.) was added. The resulting mixture was stirred at 80°C in a sealed pressure tube (Sigma-Aldrich, Cat. # Z568767) for 12 h. The solvent was then removed in vacuo and the residue was dissolved in 1 mL solution (MeOH:$H_2O$ = 1:1). The resulting mixture was filtered through a 0.22 µm PTFE membrane filter and further purified by preparative RP-HPLC. (acetonitrile/water = 1:99 to 90:10, 20 mL/min, UV 254 nm). Product fractions were combined and lyophilized to give QTZ as orange powder (12 mg, 0.03 mmol, 30%). $^1$H NMR (600 MHz, Methanol-$d_4$) δ 9.03 (d, $J$ = 4.2 Hz, 1H), 8.17 (d, $J$ = 8.9 Hz, 1H), 7.98 (d, $J$ = 8.3 Hz, 1H), 7.91 – 7.83 (m, 4H), 7.62 (s, 1H), 7.51 – 7.45 (m, 2H), 7.43 (s, 1H), 7.26 – 7.17 (m, 5H), 7.12 (s, 1H), 4.07 (s, 2H). $^{13}$C NMR (151 MHz, MeOD) δ 149.49, 130.18, 128.98, 128.67, 128.51, 128.24, 128.05, 127.60, 126.30, 125.99, 125.18, 48.00, 47.85, 47.85, 47.71, 47.62, 47.57, 47.43, 47.29, 47.17, 47.15 ESI-MS ($C_{28}H_{21}N_4O$): $[M+H]^+$ calcd: 429.16, found: 429.36.

#### 2. Luciferase Engineering and *in Vitro* and Cellular Assays

##### 2.1 Materials and General Methods

Synthetic DNA oligonucleotides were purchased from Integrated DNA Technologies. Restriction endonucleases were purchased from Thermo Fisher Scientific. High-fidelity DNA polymerase and Taq DNA polymerase were purchased from Thermo Fisher Scientific and New England Biolabs. Polymerase chain reaction (PCR) and restriction digestion products were purified by gel electrophoresis and Syd Labs Gel Extraction columns. Plasmid DNA was prepared by using Syd Labs Miniprep columns. DNA sequences were analyzed by Eurofins Genomics. HEK 293T and MDA-MB-453 cells were purchased from ATCC. The plasmids pAAV-TBG (Addgene, Cat. #

105535), pAdDeltaF6 (Addgene, Cat. # 112867) and pAAV2/9n (Addgene, Cat. # 112865) were gifts from James M. Wilson (U Penn). The plasmids pLX208-CMV (Addgene Cat. # 153007) was a gift from Alice Ting (Stanford). The plasmids pMD2.G (Addgene Cat. # 12259) and psPAX2(Addgene Cat. # 12260) were gifts from Didier Trono (EPFL).

**2.2 Library Construction by Error-Prone PCRs**

PCRs with synthetic oligonucleotide pairs pBAD_F_QLuc and pBAD_R_QLuc (Table S4) were used to amplify the LumiLuc luciferase gene from our previous pcDNA3-LumiLuc plasmid (Addgene, Cat. # 126622). Random mutations were introduced using Taq DNA polymerase with 0.2 mM $MnCl_2$ along with unbalanced dNTPs. Target luciferase genes containing random mutations were subcloned into a pBAD/His B plasmid vector predigested with Xho I and Hind III. The resultant DNA libraries were used to transform E. coli 10G competent cells (Lucigen).

**2.3 Phenotypic Screening**

E. cloni 10G cells transformed by the luciferase mutant libraries were plated on LB agar plates supplemented with ampicillin (100 μg/mL) and L-arabinose (0.1%, w/v%) and incubated at 37°C overnight to form bacterial colonies. Agar plates were left at room temperature for another 6 h, followed by BLI in a dark box (UVP Bio Spectrum) equipped with a QSI 628 cooled CCD camera (Quantum Scientific Imaging). BL images were acquired after spraying ~ 200 μL of 50 μM substrates to agar plates and further processed with the Fiji image analysis software. For each round of selection, the brightest 192 colonies from a total of ~ 20000 colonies were chosen and used to inoculate 500 μL LB broth containing ampicillin (100 μg/mL) and L-arabinose (0.1%, w/v%) in two deep 96-well plates. After overnight growth with 250 r.p.m shaking at 37 °C, the cultures were pelleted with centrifugation. The pellets were lysed with 500 μL B-PER (Thermo Fisher Scientific). 5 μL cell lysate from each well was diluted with 95 μL assay buffer (1 mM CDTA, 0.5% Tergitol NP-40, 0.05% Antifoam 204, 150 mM KCl, 100 mM MES, pH 6.0, 1 mM DTT, and 35 mM thiourea). Another 100 μL of 30 μM substrates pre-dissolved in the assay buffer were added into the lysate, resulting in the final concentration of substrates at 15 μM. BL of individual wells was measured on a CLARIOstar Microplate Reader (BMG Labtech) equipped with a reagent injector and an extended-range red-sensitive PMT. Three mutants showing the highest BL intensities or extended kinetics were chosen and subject to the next round of directed evolution.

#### 2.4 Luciferase Expression and Purification

Luciferases were expressed and purified as $His_6$-tagged fusion proteins. A single colony was picked and precultured in 5 mL LB starter culture containing ampicillin (100 μg/mL), which was next shaken at 37 °C and 250 r.p.m. overnight. The saturated overnight culture was diluted 100-fold into 2×YT broth containing ampicillin (100 μg/mL). The growth continued at 250 r.p.m. and 37 °C. When the optical density at 600 nm ($OD_{600}$) reached 0.6, the cultures were induced with the addition of L-arabinose (0.2%, w/v) and then incubated at room temperature for another 24 h. Cells were harvested by centrifugation at 4,700 r.p.m. for 15 min and lysed by sonication in the Ni-NTA wash buffer (500 mM NaCl, 20 mM Tris-HCl, pH 8.0, 30 mM imidazole). The cell debris was pelleted by centrifugation at 18,000 r.p.m. for 30 min at 4 °C, and the supernatant was loaded onto a Ni-HisTrap HP column (Cytiva), followed by washing with 20 column volume (CV) of the washing buffer. The protein was next eluted with the Ni-NTA elution buffer (500 mM NaCl, 20 mM Tris-HCl, pH 8.0, 250 mM imidazole), and then subjected to a HiLoad® 16/600 Superdex® 200 pg size-exclusion column (SEC, Cytiva) and eluted with the SEC elution buffer (30 mM Tris-HCl, 150 mM NaCl, and pH 7.4). The target-protein-containing fraction was further concentrated using 15 ml Amicon Ultra Centrifugal Filter Units with a 10-kDa cutoff (EMD Millipore). Protein concentrations were determined using the Pierce 660-nm Protein Assay Kit (Thermo Fisher) with bovine serum albumin (BSA) as the standard. Protein purity was verified using SDS-PAGE. Recombinant luciferases were aliquoted and stored at 4 °C for short-term usage and −80 °C for long-term storage.

#### 2.5 Characterization of Luciferases in *E. coli* Lysates

Three *E. coli* colonies in each group were selected and used to inoculate 3 mL LB medium containing ampicillin (100 μg/mL) and L-arabinose (0.1% w/v) at 37 °C with shaking at 250 r.p.m for 16 h. 500 μL cell cultures were taken, pelleted by centrifugation, and lysed with 500 μL B-PER. Next, 5 μL lysate was taken and further diluted with 95 μL of the assay buffer described above. 100 μL 30 μM pre-dissolved in the assay buffer were added via the reagent injector, resulting in the reaction mixture with a final concentration of the substrate at 15 μM. The spectrum was recorded by a CLARIOstar Microplate Reader (BMG Labtech) equipped with an extended-range red-sensitive PMT.

**2.6 Acquisition of Emission Spectra with Purified Luciferases**

Purified luciferases in the SEC elution buffer were diluted with the assay buffer described above to the final protein concentration of 200 nM and a total volume of 100 μL. 100 μL substrates (50 μM) pre-dissolved in the assay buffer were added to the luciferase solutions via the reagent injector. BL spectra were recorded by a CLARIOstar Microplate Reader (BMG Labtech) equipped with an extended-range red-sensitive PMT in a 5-nm interval from 400 to 800 nm.

**2.7 Enzymatic Assays with Purified Luciferases**

QLuc and NLuc were diluted to 100 pM using Dulbecco's phosphate-buffered saline (DPBS). Substrates (5 mM) dissolved in a premade solution (50% v/v ethanol, 50% v/v 1,2-propanediol, and 0.88 mg/mL L-ascorbic acid) were diluted using DPBS to gain concentrations ranging from 25 μM to 0.78 μM. 50 μL substrates were added to 50 μL of the luciferase DPBS solution using a multichannel pipette. BL intensities were recorded by a CLARIOstar Microplate Reader (BMG Labtech) equipped with an extended-range red-sensitive PMT. The experiments were repeated three times, and the initial brightness was recorded for calculating $K_M$ and relative $k_{cat}$. Data were fitted in GraphPad Prism 8 using the Michaelis-Menten nonlinear regression function.

**2.8 Plasmid Construction and Characterization in HEK 293T cells**

pBAD-QLuc was used as the template for PCR with oligos pcDNA3_F_QLuc and pcDNA3_R_QLuc (Table S4). The resulting DNA fragment was inserted into a pcDNA3 plasmid predigested with Hind III and Xho I to afford the plasmid pcDNA3-QLuc. An extra start codon (ATG) was introduced before the QLuc sequences for transcription initiation. HEK 293T cells were cultured and transfected with 3 μg of the pcDNA3 plasmid as previously described.[1] Cell numbers were quantified using a hemocytometer. 10,000 transiently transfected cells were diluted with DPBS to a total volume of 100 μL and placed in individual wells in a 96-well plate. The corresponding 100 μL of 100 μM luciferin substrates in DPBS were added to get a final concentration of 50 μM substrates. BL images were acquired in a dark box (UVP Bio Spectrum) equipped with a Computer Motorized ZOOM lens (M6Z1212MP3) and an Andor iXon Life 888 EMCCD camera. The Andor iXon Life 888 EMCCD was first set to the "Photon Counting" mode using the pre-defined OptAcquire options, but the gain was reduced from 750 to 2. The exposure time was 5 s with 1×1 binning, and the acquisition intervals were 30 s. The ZOOM lens was set to 100% open aperture, 0% zoom, and 0% focus using the UVP VisionWorksLS software. The plate was placed 21 cm away from the front of the lens. A 600-700 nm filter (Chroma, Cat. # 27014) was placed in front of the camera lens during imaging acquisition. All images were processed and pseudo-colored using the Fiji image analysis software.

#### 3. In Vivo BLI and Relevant Experiments

##### 3.1 AAV Packing and Quantification of Viral Titers

The NLuc gene was amplified from pcDNA3-NLuc using oligos pAAV_F_NLuc and pAAV_R_NLuc (Table S4), then cloned into a pAAV-TBG vector (Addgene, Cat. # 105535) between Nco I and BamH I to generate pAAV-TBG-NLuc. pAAV-TBG-teLuc and pAAV-TBG-QLuc were similarly constructed. Next, the pAAV plasmids along with pAdDeltaF6 (Addgene, Cat. # 112867) and pAAV2/9n (Addgene, Cat. # 112865), were used to transfect HEK 293T cells. A protocol by Rego et al.[2] was followed for virus packing and purification. Viral titers were quantified with quantitative PCR (qPCR) by following a protocol from Addgene. Typically, AAV titers ranging from $1\times10^{13}$ - $1\times10^{14}$ GC/mL were acquired, AAVs were aliquoted and stored at −80 °C for long-term stability.

##### 3.2 Comparison of Luciferases in the Liver in Live Mice.

AAVs were diluted to the desired concentration ($5\times10^{12}$ GC/mL) using DPBS before use. 100 µL of the virus was systematically delivered to BALB/cJ mice (The Jackson Laboratory, Cat #000615) via tail vein injection. Three weeks later, mice were utilized for BLI. 100 µL of 1 mM luciferins dissolved in the in vivo injection buffer (25% (w/v) 2-hydroxypropyl-β-cyclodextrin (HP-β-CD) and 20% (v/v) PEG-400 dissolved in normal saline) were delivered into mice intraperitoneally. BLI was performed in the aforementioned BLI dark box immediately after substrate administration. The Andor iXon Life 888 EMCCD was first set to the "Photon Counting" mode using the pre-defined OptAcquire options. The gain was reduced to 500. Camera binning was set at 1×1, and camera temperature was set at -70 °C. Exposure time was 5 s, and acquisition interval was 30 s. The ZOOM lens was set to 100% open aperture, 0% zoom, and 0% focus set to 0% using the UVP VisionWorksLS software. For each experiment, three anesthetized mice expressing different luciferases were placed side-by-side and 21 cm away from the front of the lens. Images were processed and quantified using the Fiji image analysis software. Data were plotted and statistical comparison was performed in GraphPad Prism 8.

##### 3.3 Plasmid Construction and Expression of the scFv-QLuc Fusion Protein

PCR with synthetic oligonucleotide pairs pET_F1_scFvQLuc and pET_R1_scFvQLuc (Table S4) was used to amplify the scFv gene from a custom synthetic gene fragment (Integrated DNA Technologies). The synthetic oligonucleotide pairs pET_F2_scFvQLuc and pET_R2_scFvQLuc were used to amplify the QLuc gene from the bacterial expression plasmid. Next, the amplified scFv and QLuc fragments were used for three-fragments Gibson assembly along with the pET28a plasmid predigested with Nco I and Xho I. The resulting plasmid was used to transform SHuffle T7 Express Competent Cells (NEB, Cat # C3029J), which was reported to benefit the expression and folding of scFvs.[3] Colonies were formed after overnight culture on 2×YT agar plates containing 50 μg/mL kanamycin. A single fresh colony was picked for preculturing in 5 mL 2×YT broth containing 50 μg/mL kanamycin. After overnight growth at 37 °C and 250 r.p.m., the preculture was diluted 100-fold with 500 mL 2×YT broth containing 50 μg/mL kanamycin. When $OD_{600}$ reached 0.6, IPTG at the final concentration of 1 mM was used to induce protein expression. The culture was shaken at 16 °C and 250 r.p.m. for 2 days. The scFv-QLuc fusion protein was lysed by sonication and purified using Ni-NTA affinity chromatography and SEC column chromatography.

##### 3.3 Immunoassays to Verify the Binding of scFv-QLuc to Her2

HeLa and MDA-MB-453 cells were trypsinized and resuspended in 5 mL DMEM. Cell numbers were determined using a hemocytometer and diluted to $1\times10^5$ cells/mL in DPBS. 1 mg of the purified scFV-QLuc fusion protein was added to 1 mL of the cell suspension, which was next incubated on ice for 30 minutes. Cells were then gently pelleted by centrifugation and resuspended in 1 mL DPBS. The centrifugation and resuspension procedures were conducted three times. Next, 100 µL of cell suspension was added to a 96-wells plate. 100 µL of 50 µM QTZ dissolved in DPBS was added to the cell suspension via a reagent injector, resulting in the final concentrations of QTZ at 25 µM. BL intensities were recorded on a CLARIOstar Microplate Reader (BMG Labtech) equipped with an extended-range red-sensitive PMT. The endpoint reading mode was used, and these experiments were repeated three times.

##### 3.4 Lentivirus Packing and Establishment of Akaluc-Expressing Stable Cell Lines

The Akaluc gene was amplified using oligos Plx208-F-Akluc and Plx208-R-Akluc (Table S4) and inserted into a pLX208-CMV plasmid (Addgene Cat. # 153007) vector between restriction sites

BamH I and Mlu I. HEK 293T cells at ~ 70% confluency were co-transfected with pLX208-CMV-AkaLuc and lentiviral packing plasmids pMD2.G (Addgene Cat. # 12259) and psPAX2(Addgene Cat. # 12260). Cell culture media were replaced with fresh media 6-8 h post-transfection, and cells were cultured for an additional 48 h. Next, the cell culture media containing lentiviruses were collected, filtered through 0.45 μm PES syringe filters (Denville Scientific, Cat. # 1159T84), and concentrated using a PEG-8000 based precipitation method from the MD Anderson Functional Genomics Core. HEK 293T cells or MDA-MB-453 cells at ~70% confluency were transduced with the lentivirus. Three days later, fresh media containing 50 μg/mL hygromycin were used to culture the cells, and cells were maintained in this condition for another 5 days. Survived cells were propagated and maintained as polyclonal stable cell lines in the presence of 100 μg/mL hygromycin.

**3.5 Establishment of a Xenograft Tumor model with Akaluc-Labeled Cells**

HeLa and MDA-MB-453 cells stably expressing Akaluc luciferases were trypsinized and resuspended in 5 mL DMEM. Cell numbers were determined using a hemocytometer, and cell viability was determined using a trypan blue exclusion test. $1 \times 10^6$ HeLa cells or $3 \times 10^6$ MDA-MB-453 cells were resuspended in a pre-mixture of 50 μL DMEM and 50 μL Matrigel basement membrane matrix (Corning, Cat. # 356237). Four-week-old female NU/J mice (The Jackson Laboratory, Cat #002019) were anesthetized using ketamine. Cells were subcutaneously injected to the left and right shoulders in mice.

**3.6 In Vivo ImmunoBLI of Her2(+) Tumors**

Three weeks after tumor implantation, mice were utilized for BLI. Mice were first anesthetized using ketamine and xylazine. The scFv-QLuc fusion protein (1 mg) diluted with normal saline to a volume of 100 μL was administered through the tail vein. 6 h later, 100 μL QTZ (1 mM) dissolved in the in vivo injection buffer mentioned above was also administered through intravenous injection. BLI was performed immediately after substrates administration in the aforementioned BLI dark box. The instrumental settings were identical to those described under Section 3.2, except that 10-s exposure with a 1-min acquisition interval was used. Anesthetized mice were placed 16 cm away from the front of the lens. After imaging, mice recovered from anesthesia were returned to the vivarium. During the next day, when residual signals from QLuc-QTZ were dissipated, 100 μL Akalumine (1 mM) dissolved in the aforementioned in vivo injection buffer was administered intraperitoneally. Imaging was performed with identical instrumental settings immediately after substrates administration. Data were processed and quantified using the Fiji image analysis software. Data were plotted and statistical comparison was performed in GraphPad Prism 8.

### NMR Spectra

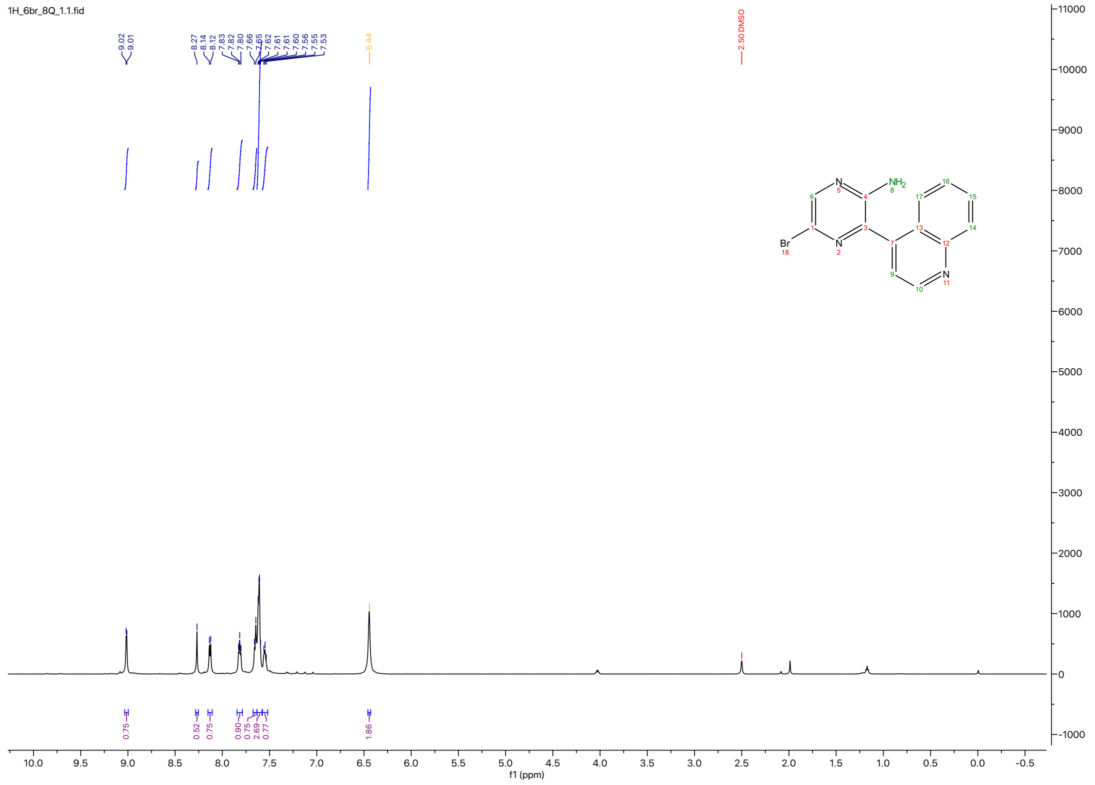


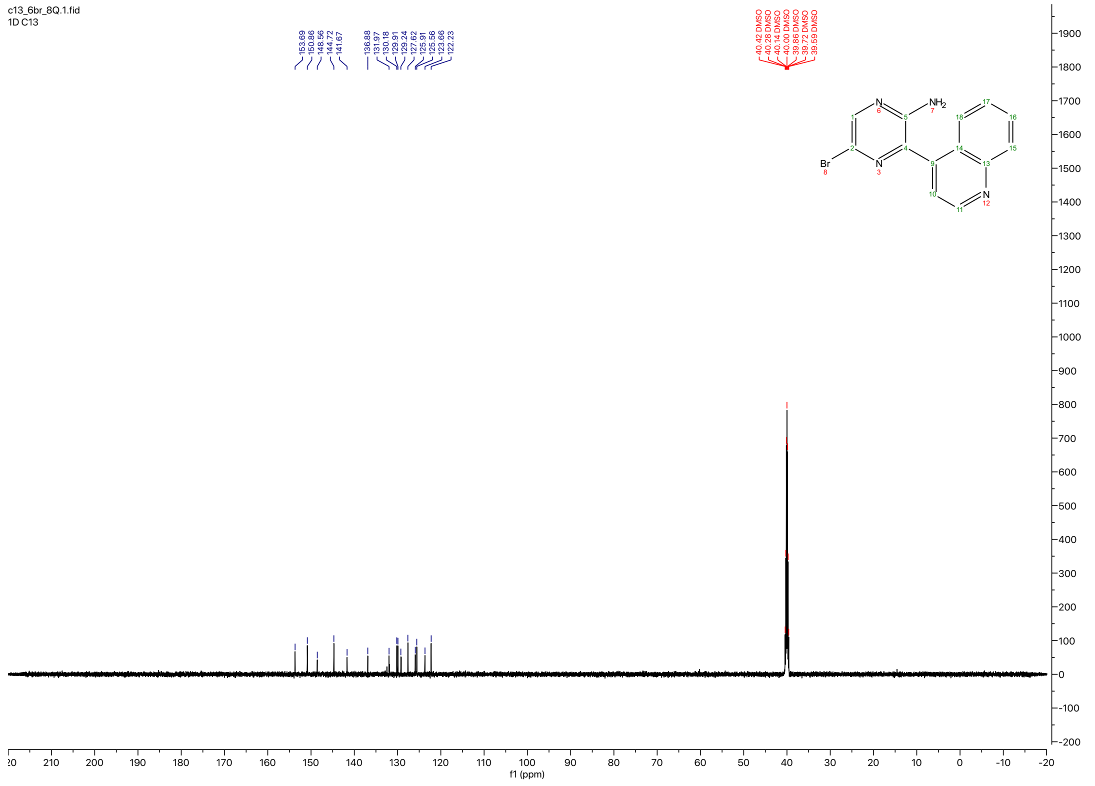

1H_6phe_8Q_1.1.fid

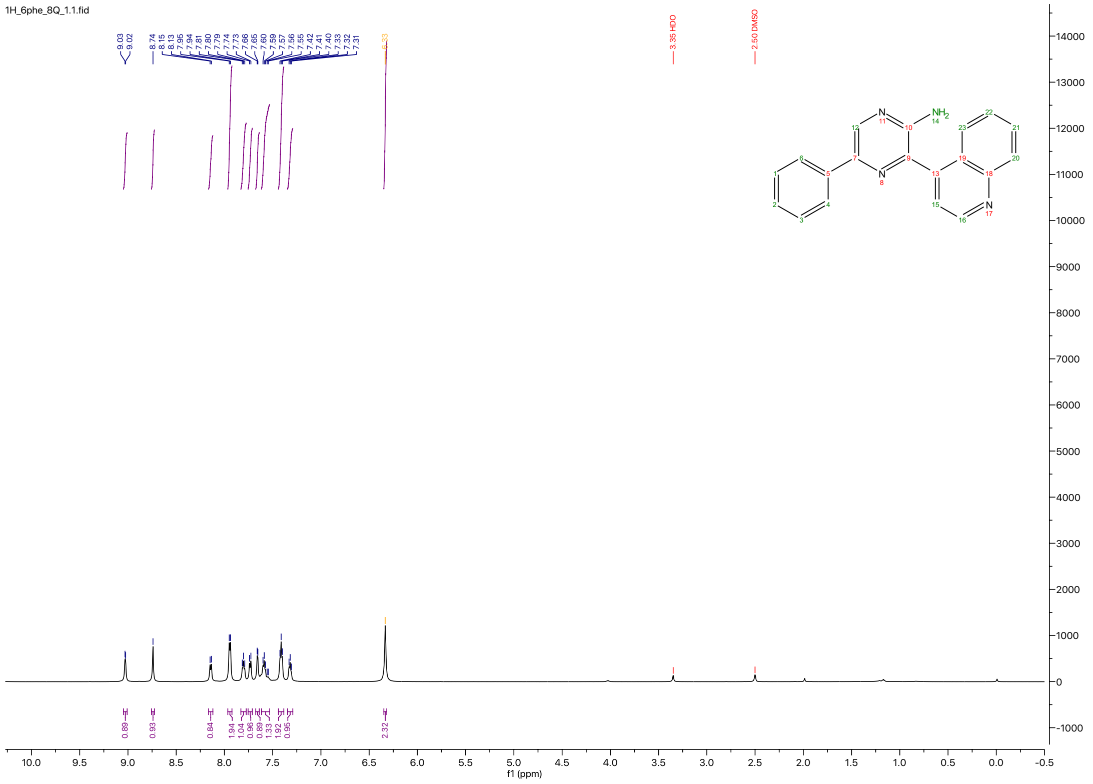

13C_6phe_8Q.1.fid
1D C13

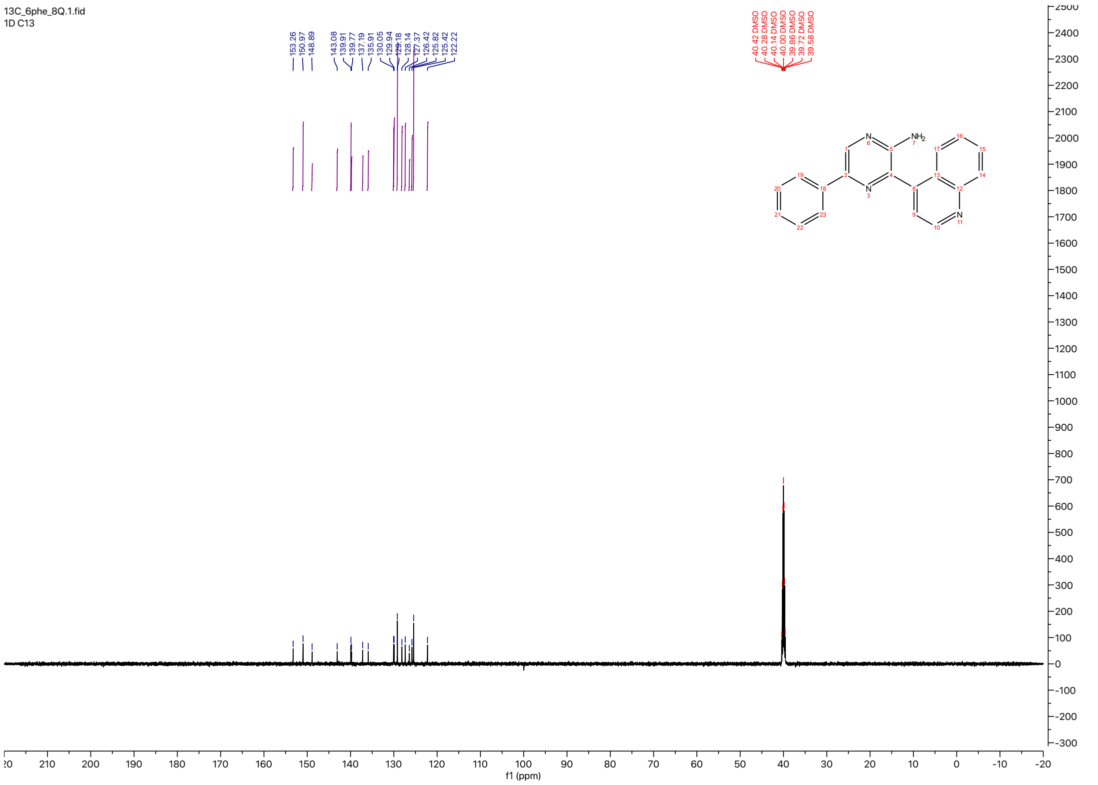

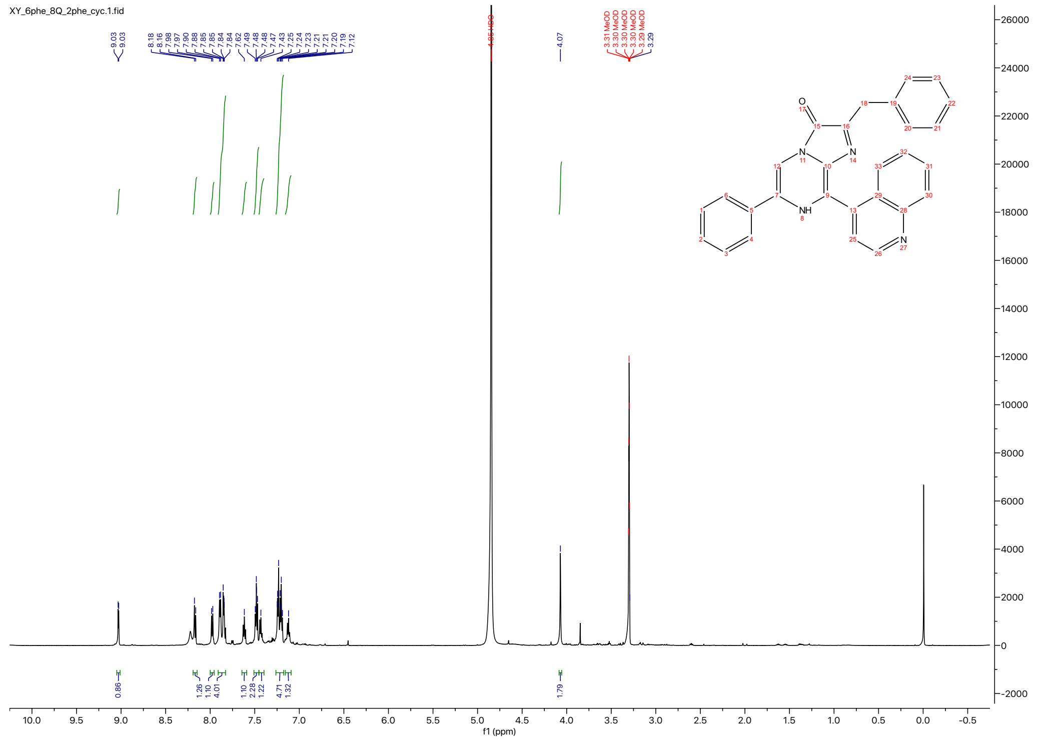
XY_6phe_8Q_2phe_cyc.1.fid
f1 (ppm)

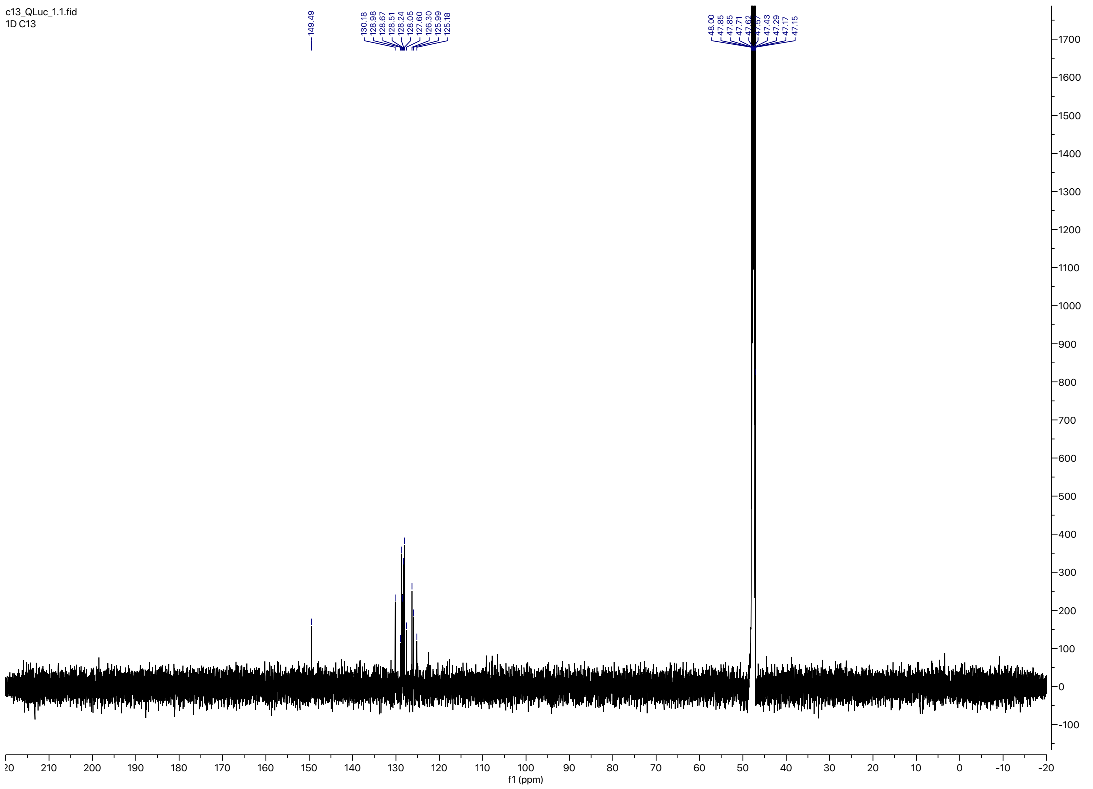
c13_QLuc_1.1.fid
1D C13
f1 (ppm)
